# Structure of the molecular bushing of the bacterial flagellar motor

**DOI:** 10.1101/2020.11.12.379347

**Authors:** Tomoko Yamaguchi, Fumiaki Makino, Tomoko Miyata, Tohru Minamino, Takayuki Kato, Keiichi Namba

**Author notes:** These authors contributed equally: Tomoko Yamaguchi, Fumiaki Makino. Correspondence and requests for materials should be addressed to K.N. or T.K.

## Abstract

The bacterial flagellum is a motility organelle, consisting of the basal body acting as a rotary motor, the filament as a helical propeller and the hook connecting these two as a universal joint^1,2^. The basal body contains three rings: the MS ring as the transmembrane core of the rotor; the C ring essential for torque generation and switching regulation; and the LP ring as a bushing supporting the distal rod for its rapid, stable rotation without much friction. The negatively charged surface of the distal rod suggested electrostatic repulsive force in supporting high-speed rotation of the rod as a drive shaft^3^, but the LP ring structure was needed to see the actual mechanisms of its bushing function and assembly against the repulsive force. Here we report the LP ring structure by electron cryomicroscopy at 3.5 Å resolution, showing 26-fold rotational symmetry and intricate intersubunit interactions of each subunit with up to six partners that explains the structural stability. The inner surface is charged both positively and negatively, and positive charges on the P ring presumably play important roles in its initial assembly around the rod in the peptidoglycan layer followed by the L ring assembly in the outer membrane.

---

The bacterial flagellum is a motility organelle responsible for rapid movement of bacterial cells towards more desirable environments. The flagellum consists of three structural parts: the basal body working as a rotary motor, the filament as a screw propeller, and the hook as a universal joint connecting the filament to the motor. The flagellar motor converts the electrochemical potential difference of cations across the cell membrane to mechanical work required for high-speed rotation with almost 100% efficiency^4^, and the maximum rotation speed has been measured to be 1,700 revolutions per second (rps)^5^, which is much faster than that of the Formula One racing car engine.

The basal body consists of the C ring, MS ring, LP ring and rod. The MS-C ring acts as a rotor of the flagellar motor. The MS ring is the transmembrane core of the rotor and the initial template for flagellar assembly. The C ring is attached to the cytoplasmic face of the MS ring and is essential for torque generation and switching regulation of the motor. The rod is a rigid, straight cylindrical structure directly connected with the MS ring and acts as a drive shaft that transmits the motor torque to the filament through the hook. It is divided into two structural parts: the proximal rod composed of FliE, FlgB, FlgC and FlgF; and the distal rod formed by FlgG. The LP ring is a bushing supporting the distal rod for its rapid, stable rotation without much friction and is composed of a lipoprotein, FlgH, and a periplasmic protein, FlgI. The L and P rings are embedded within the outer membrane and the peptidoglycan (PG) layer, respectively^1,2^ (Fig. 1a). The LP ring is very stable against various chemical treatment^6,7^, and its inner surface must be smooth to sustain high-speed rotation of the rod even up to around 1,700 rps^5^. High-resolution structures of the distal rod by electron cryomicroscopy (cryoEM) and X-ray crystallography have revealed its outer surface being highly negatively charged^3,8^, leading to a plausible hypothesis that the inner surface of the LP ring may also be negatively charged to generate electrostatic repulsive force to keep the rod rotating at the center of the LP ring to minimize friction between them. However, how the LP ring assemble around the rod against the repulsive force remains unclear.

**Fig. 1.**
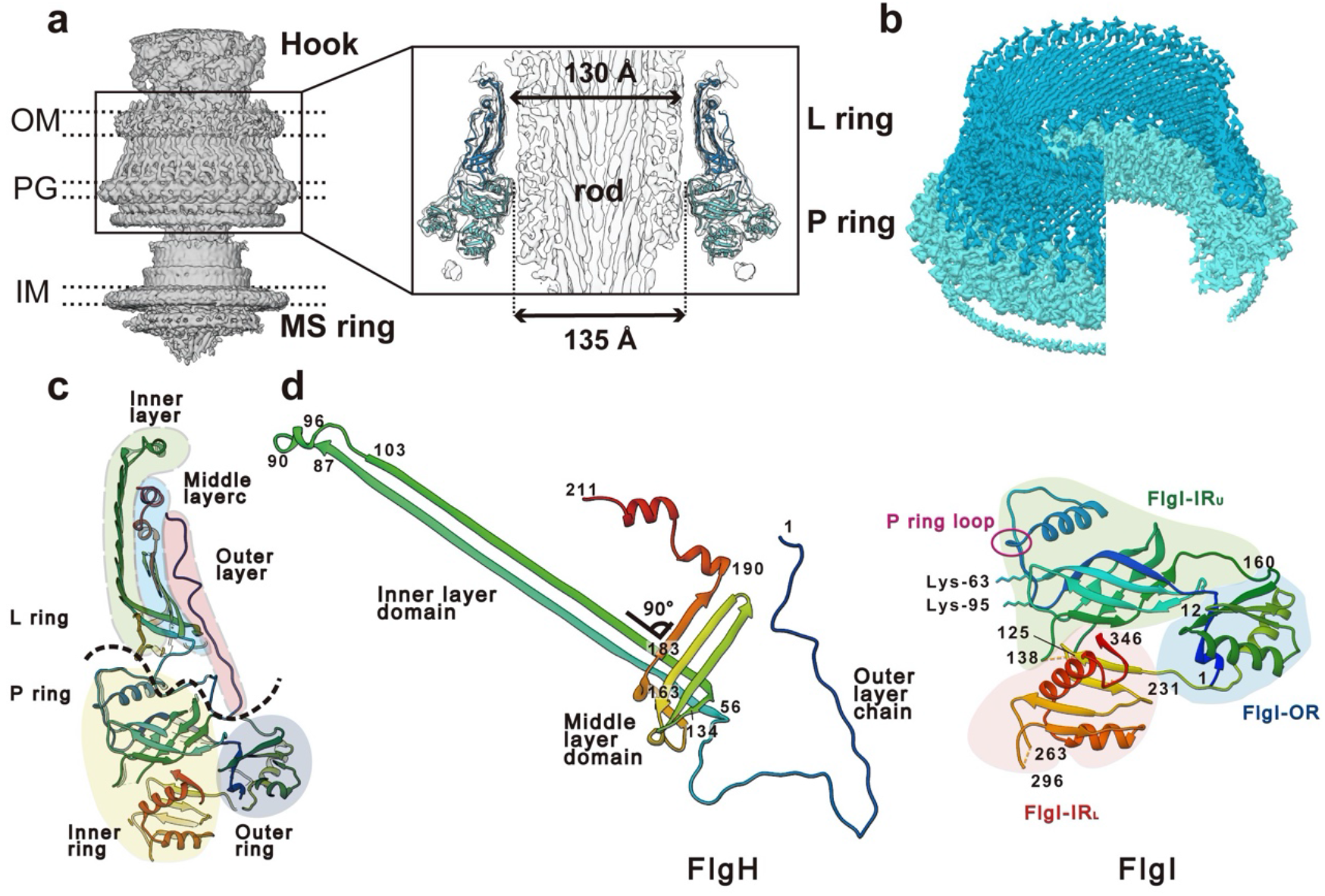
Density map and the atomic model of the LP ring. **a**, Side view of the density map of the HBB complex (EMD-30409) without applying any symmetry (C1). OM, PG, IM refers to the outer membrane, peptidoglycan layer and inner membrane, respectively. A vertical slice of the LP ring and rod is shown enlarged on the right. The atomic models of the L ring (sky-blue) and the P ring (cyan) are superimposed to the density map. **b**, the density maps of the L ring (sky-blue) and the P ring (cyan) after applying C26 symmetry (EMD-30398). **c**, The atomic model of FlgH and FlgI in Cα ribbon representation in a vertical slice of the LP ring. The L ring has a three-layered structure (inner layer, light green; middle layer, cyan; outer layer, pink), and the P ring has a two-layered structure (inner ring, beige; outer ring, blue). **d**, The atomic model of FlgH and FlgI (PDB ID: 7CLR) in Cα ribbon representation. FlgH is constituted from three parts: the inner layer domain; the middle layer domain; the outer layer chain. FlgI is constituted from three domains: FlgI-OR (cyan); FlgI-IR_U_ (light green); and FlgI-IR_L_ (pink). Lys-63, Lys-95 and the P ring loop are located within the FlgI-IR_U_ domain.

The flagellar axial proteins for assembly of the rod, hook and filament are transported from the cytoplasm to the distal end of their growing structures by a specialized protein export apparatus within the MS-C ring. In contrast, FlgH and FlgI are synthesized as precursors with a cleavable N-terminal signal peptide and are translocated to the periplasm by the Sec translocon^9^. The N-terminal signal peptides are cleaved during the translocation across the cell membrane^10,11^. FlgI molecules assemble around the distal rod with the help of the FlgA chaperone to form the P ring^12^. The P ring tightly associates with the PG layer, allowing the LP ring to act as a molecular bushing. The N-terminal domain of FlgI consisting of residues 1–120 is critical not only for the stabilization of the P ring structure but also for the formation of the LP ring complex^13^. Because the L ring is not formed in *flgI* null mutants^7^, FlgH assembly around the rod presumably requires an interaction between FlgH and FlgI. Certain point mutations in the distal rod protein FlgG not only produces abnormally elongated rods called the polyrod but also allow many P rings to be formed around them, suggesting that the well-regulated P ring formation involves interactions of specific amino acid residues of FlgG and FlgI^14,15^. However, the detailed mechanism of LP ring formation remains elusive due to the lack of structural information.

To clarify the interactions between the rod and LP ring as well as the assembly mechanism of the LP ring, we carried out cryoEM structural analysis of the LP ring in the flagellar basal body from S*almonella enterica* serovar Typhimurium (hereafter *Salmonella*). Here we report the LP ring structure at 3.5 Å resolution, showing 26-fold rotational symmetry and intricate intersubunit interactions of each FlgH subunit with six FlgH partners that explains the structural stability. The inner surface is charged both positively and negatively where positive charges on the P ring appear to play important roles in its initial assembly around the rod.

## Structure of the LP ring around the rod

We analyzed the LP ring structure in the hook-basal body (HBB) by cryoEM single particle image analysis. First, the three-dimensional (3D) image of the HBB was reconstructed at 6.9 Å resolution with C1 symmetry from 13,017 HBB images extracted from 12,759 cryoEM movie images (Extended Data Figs. 1 and 2). This density map contains the MS ring, LP ring, rod and proximal end of the hook. The C26 symmetry was visible in the P ring density and was confirmed by autocorrelation along the circumference (Extended Data Fig. 2b; see Methods for mor detail). In order to obtain a higher resolution structure of the LP ring, the LP ring part of the HBB image was re-extracted and analyzed with C26 symmetry, which produced a 3.5 Å resolution map (EMD-30398). To determine the relative positioning of the rod and LP ring, another 3D image of the HBB was reconstructed at 6.9 Å resolution with C1 symmetry from 14,370 HBB images (Fig. 1a, Extended Data Fig. 2a, EMD-30409). The inner and outer diameters of the LP ring are 135 Å and 260 Å, respectively, and its height is 145 Å (Fig. 1a). The inner diameter of the LP ring is only slightly larger than the diameter of the rod (130 Å). There is an extra, rather blurred ring density beneath the P ring, which is likely part of FlgI (Fig. 1b). The position of the LP ring around the rod was determined by superimposing this high-resolution map on the HBB density map (Extended Data Fig. 3).

## Structures of FlgH and FlgI in the LP ring

We built the atomic models of FlgH and FlgI based on the density map (PDB ID: 7CLR). The atomic model of FlgH forming the L ring contains residues 1–211. The L ring adopts a three-layer structure composed of two β-barrel layers and the outermost layer with an extended chain (Fig. 1c,d, Extended Data Fig. 4). Two very long anti-parallel β strands, a short β strand and a short α helix make the inner layer domain, four anti-parallel β strands and two short α helices make the middle layer domain, and the N-terminal extended chain (outer layer chain) covers these two layers to form the outer layer. The anti-parallel β strands of the inner and middle layer domains are crossing nearly perpendicular to each other (Fig. 1d), forming a hydrophobic core with conserved residues (Extended Data Fig. 4a,c), and 26 copies of them form two layers of cylindrical β sheets in the inner and middle layers, respectively (Fig. 1c and 2a,b). The cylindrical β barrel structure is similar to those of secretin proteins, such as GspD of the type II secretion system^16^ and InvG of the type III secretion system^17^, except for the orientation of the long β strands and the presence of the extended chain in the outer layer of the L ring (Extended Data Fig. 6). As the outer layer chain of FlgH is predicted to be disordered (Extended Data Fig. 4b), this part is presumably disordered in the monomeric form of FlgH until it assembles into the L ring.

**Fig. 2.**
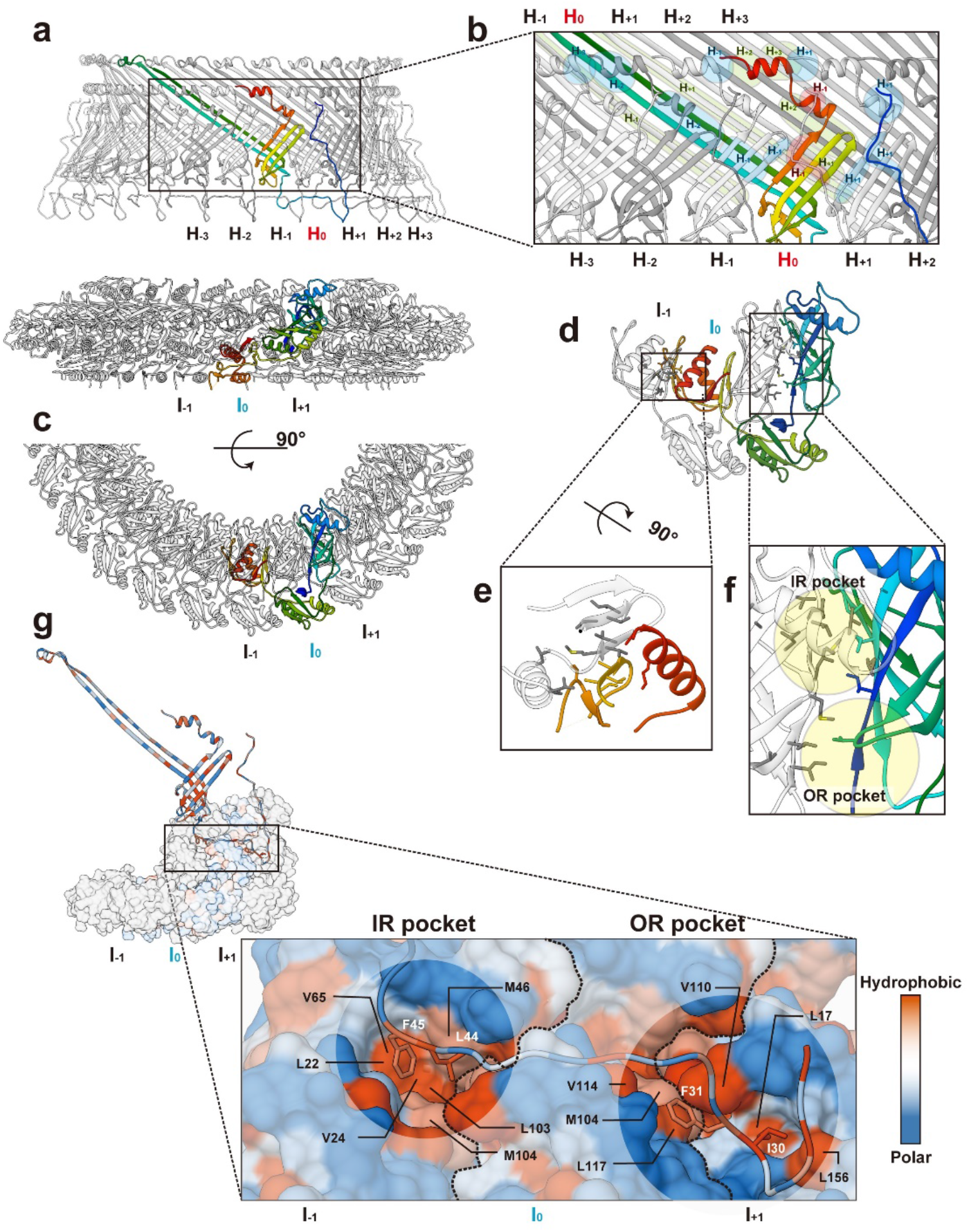
Intersubunit interactions of FlgH and FlgI in the LP ring. **a**, Side view of the L (upper panel) and P (lower panel) rings with one subunit colored rainbow. **b**, Intersubunit interactions between a FlgH subunit (H_0_) and its neighboring six FlgH subunits (H_±1_, H_±2_, H_±3_). The segments of six neighboring FlgH subunits interacting with the subunit H_0_ are labelled with their subunit ID and colored as: light green, interactions with the inner layer; light blue, interactions with the middle layer, pink, interactions with the outer layer. **c**, **d**, Top view of the P ring, showing the intersubunit interactions between FlgI (I_0_) and its neighboring subunit (I_±1_). **e**, Hydrophobic interactions and a short anti-parallel β sheet formed by two neighboring FlgI-IR_L_ domains. **f**, Two hydrophobic pockets (IR and OR pockets) formed by two neighboring FlgI-IR_U_ domains. Hydrophobic side chains are displayed in grey sticks in **e**and **f**. **g**, Interactions between FlgH (H_0_) and three FlgI (I_0_, I_±1_) subunits. The extended chain of FlgH in the outer layer form hydrophobic interactions with the IR and OR pockets formed by FlgI subunits.

The atomic model of FlgI contains residue 1–125, 138–263 and 296–346 (Fig. 1d, Extended Data Fig. 5). It is composed of three domains, which we named FlgI-IRU (upper inner ring), FlgI-IRL (lower inner ring) and FlgI-OR (outer ring). FlgI-OR (residues 1–11 and 160–230) is composed of three α helices and four β strands and forms the outer rim of the P ring. FlgI-IRU (residues 12–159) is composed of one α helix and 10 β strands, FlgI-IRL (residues 231–346) is composed of two α helices and five β strands, and these two domains form the inner part of the P ring. The structure of FlgI-OR looks similar to that of the N3 domain of GspD (residues 232-267, 288-323)^16^, the N3 domain of InvG (residues 176-227, 252-302)^17^, the N1 domain of PilQ (residues 574-639), which is an outer membrane protein of type VIa pili^18^, and the C-terminal domain of MotY (residues 155–202, 210-243, 262-271)^19^, which shows an extensive similarity to the PG binding domain of the OmpA family protein interacting with the PG layer (Extended Data Fig. 6 and 7; analyzed by MATRAS^20^). Therefore, FlgI-OR is likely to interact with the PG layer directly. FlgI-IRU forms a β-barrel-like structure with highly conserved residues forming a hydrophobic core (Fig. 1d, Extended Data Fig. 5a,b) and also contains a loop of four polar residues from Thr-33 through Thr-36 (P ring loop in Fig. 1d). This loop and two highly conserved positively charged residues of FlgI-IRU, Lys-63 and Lys-95, are in very close proximity to the rod surface, suggesting their involvement in P ring assembly around the rod as well as in the function as a bushing. FlgI-IRL is formed by the C-terminal chain of FlgI and contains a highly flexible loop (residues 264-295) not visible in the density map.

## Intersubunit interactions of FlgH and FlgI in the LP ring

In the L ring structure, each FlgH subunit (H_0_) interacts with six adjacent FlgH subunits (H_±1_, H_±2_ and H_±3_) (Fig. 2a,b), and these intricate intersubunit interactions make the LP ring mechanically stable. In the P ring, each FlgI subunit (I_0_) interacts with two adjacent FlgI subunits (I_−1_ and I_+1_) (Fig. 2a,c,d). The FlgI-IR_L_ domains of subunits I_0_ and I_−1_ interact with each other through hydrophobic interactions and also form a short anti-parallel β sheet (Fig. 2e). FlgI-IR_U_ forms two hydrophobic “pockets” named the IR (inner ring) and OR (outer ring) pockets (Fig. 2f). The C-terminal half of the outer layer chain of a FlgH subunit (H_0_) lies at the bottom of the L ring and interacts with three FlgI subunits (I_0_, I_±1_) through its interactions with the hydrophobic IR and OR pockets of the P ring (Fig. 2g). Leu-44 and Phe-45 of FlgH are in the IR pocket composed of Leu-22, Val-24, Met-46, Val-65, Leu-103 and Met-104 of FlgI (Fig. 2g left). In the OR pocket, Ile-30 of FlgH interacts with Leu-17 and Leu-156 of FlgI, and Phe-31 of FlgH interacts with Met-104, Val-110, Val-114 and Leu-117 of FlgI (Fig. 2g right). The residues of these hydrophobic interactions, especially those in the IR pocket, are well conserved (Extended Data Fig. 5c and 9b,c).

## Surface potential and conservation of the LP ring and rod

In contrast to the prediction that electrostatic repulsive force by the negative charges on the surfaces of the rod and LP ring may play an important role in the bushing function, the inner surface of the LP ring is charged both negatively and positively. Asp-78, Asp-86 and Glu-104 of FlgH form a negative charge belt, and Lys-69 and Lys-114 of FlgH form a positive charge belt, suggesting that both repulsive and attractive force keep the negatively charged rod rotating at the center of the L ring (Fig. 3a). We also found that Lys-63 and Lys-95, which are relatively well conserved among FlgI homologues, form a positive belt on the inner surface of the P ring (Fig. 3a, Extended Data Fig. 9a), and these residues are in very close proximity to a negative charge cluster on the rod outer surface, raising a plausible hypothesis that these two lysine residues are critical for P ring assembly around the rod.

**Fig. 3.**
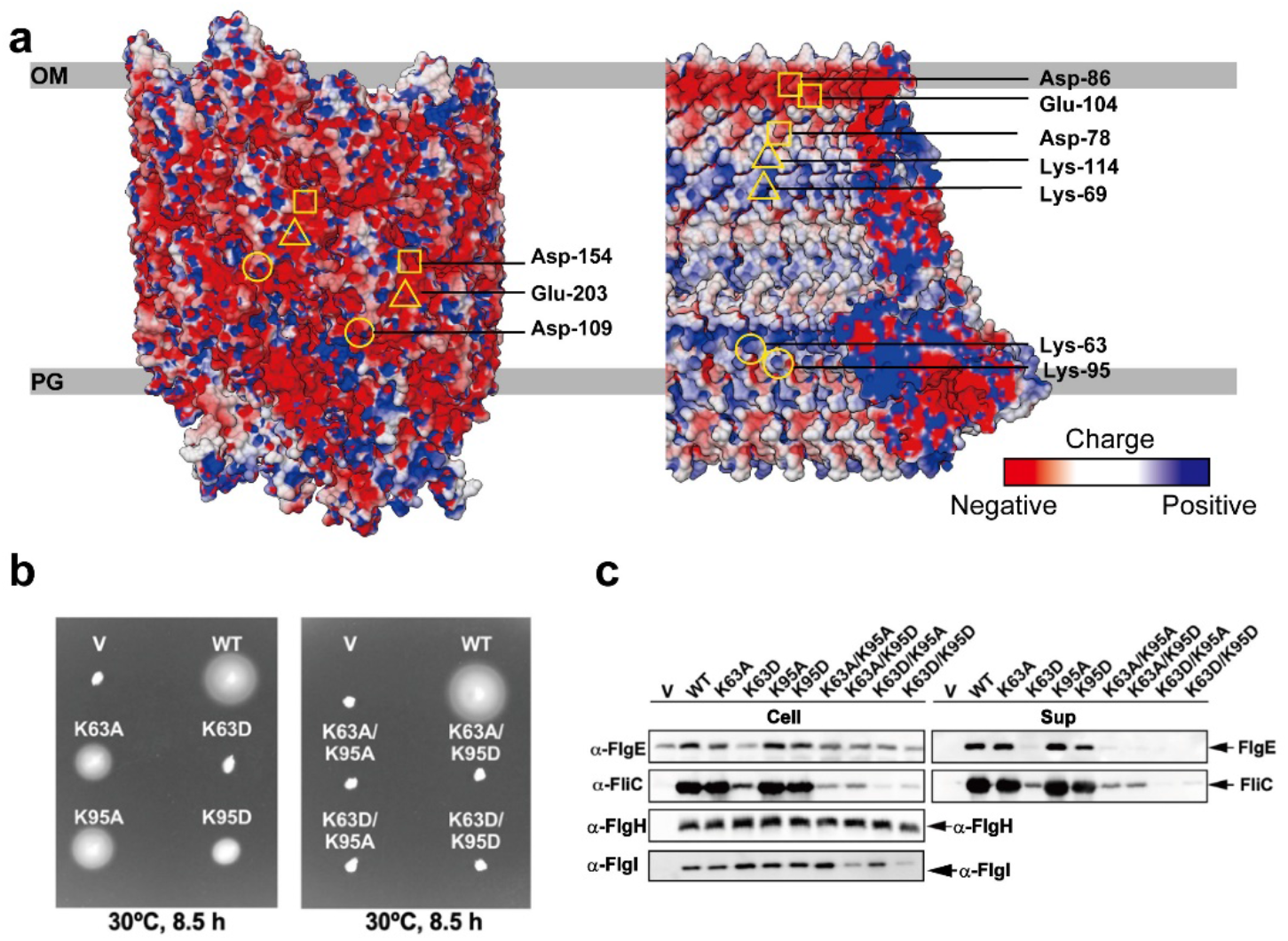
Surface potential of the rod and LP ring with *flgI* mutation analysis. **a**, Electrostatic potential of the outer surface of the distal rod and the inner surface of the LP ring. The helical arrangement of Asp-109 (○), Asp-154 (□) and Glu-203 (△) of FlgG on the surface of the rod form negatively charged belts. Asp-86, Glu-104 and Asp-78 of FlgH (□) form a negatively charged belt on the inner surface of the L ring. Lys-69 and Lys-114 (△) of FlgH and Lys-63 and Lys-95 (○) of FlgI form positively charged belts on the inner surface of the LP ring. **b**, Swarm motility assay on the soft agar plate (0.35%) of *flgI(K63A)*, *flgI(K63D) flgI(K95A) flgI(K95D)* (left) and *flgI(K63A/K95A)*, *flgI(K63A/K95D)*, *flgI(K63D/K95A)* and *flgI(K63D/K95D)* (right). **c**, The cellular expression levels of FlgE, FliC, FlgH and FlgI (left) and the secretion levels of FlgE and FliC (right) by immunoblotting with polyclonal anti-FlgE, FliC, FlgH and FlgI antibodies.

To examine this hypothesis, we constructed eight *flgI* mutants, *flgI*(K63A), *flgI*(K63D), *flgI*(K95A), *flgI*(K95D), *flgI*(K63A/K95A), *flgI*(K63A/K95D), *flgI*(K63D/K95A) and *flgI*(K63D/K95D), and analyzed their motility on 0.35% soft agar (Fig. 3b). The *flgI*(K63A), *flgI*(K95A) and *flgI*(K95D) mutants formed motility rings although not at the wild-type level, but the *flgI*(K63D), *flgI*(K63A/K95A), *flgI*(K63A/K95D), *flgI*(K63D/K95A) and *flgI*(K63D/K95D) mutants exhibited no motility. The *flgI*(K63A/K95D) and *flgI*(K63D/K95D) mutation affected the protein stability of FlgI, explaining their no motility phenotype, but other mutations did not affect protein stability at all (Fig. 3c), indicating that both Lys-63 and Lys-95 are critical for the FlgI function.

The LP ring is required for the secretion of export substrates, such as the hook capping protein FlgD, the hook protein FlgE and the filament protein FliC, into the culture media^21^. In the absence of the LP ring, FlgD and FlgE can cross the cytoplasmic membrane but not the outer membrane, suggesting that the LP ring is required for FlgD cap assembly at the distal end of the rod to initiate hook assembly by FlgE outside the cell body. To test whether these FlgI mutations affect P ring assembly, we analyzed the secretion levels of FlgE and FliC by immunoblotting with polyclonal anti-FlgE and anti-FliC antibodies (Fig. 3c). The FlgE and FliC secretion by the *flgI*(K63A) and *flgI*(K95A) mutants were at the wild-type level. Because the swarm plate motility of these two mutants was about 2-fold lower than that of the wild-type, the K63A and K95A mutations probably reduced the rate of P ring assembly by 2-fold. The FlgE and FliC secretion by the *flgI*(K95D) mutant was lower than the wild-type level, suggesting that this mutation significantly affects P ring assembly. The K63D, K63A/K95A, K63A/K95D, K63D/K95A and K63D/K95D mutations considerably reduced the secretion levels, suggesting that these five mutations inhibit P ring formation. We therefore propose that electrostatic interactions of Lys-63 and Lys-95 of FlgI with a negative charge cluster on the surface of the rod are required for P ring formation around the rod.

## Discussion

The atomic model of the LP ring around the rod provided many insights into the mechanical stability, bushing function, and assembly mechanism of the LP ring. The LP ring is stable even against a harsh treatment by acids and urea^6,7^. This extremely high stability can now be explained by the intricate intersubunit interactions between FlgH subunits, between FlgI subunits and between FlgH and FlgI. FlgH forms the three-layer cylinder structure of the L ring, consisting of two cylindrical β-barrels with their β-strands oriented nearly perpendicular to each other and the extended chain in the third layer lining them on the outer surface (Fig. 1c and 2a,b). Each FlgH subunit interacts with six adjacent FlgH subunits, making the L ring structure even more stable. Intersubunit interactions of FlgI in the P ring are not so extensive as those of FlgH in the L ring but each of FlgI three domains intimately interacts with domains of the adjacent subunits, even by forming intersubunit β-sheets (Fig. 2c-f). Then the extensive hydrophobic interactions between the extended chain of FlgH at the bottom surface of the L ring and the two pockets of FlgI on the upper surface of the P ring make the entire LP ring structure very stable (Fig. 2g).

The outer surface of the rod and inner surface of the LP ring are both very smooth and form a small gap between them that is large enough to accommodate one or two layers of water molecules in most part of their interfaces (Fig. 1a), indicating that these two structures are optimally designed for free rotation of the rod inside the LP ring without much friction. The outer surface of the rod is highly negatively charged^3,8^ but the inner surface of the LP ring is charged both negatively and positively, probably to balance repulsive and attractive forces to keep the rod stably positioned and rotating at the center of the LP ring (Fig. 3a and 4a-e). Nanophotometric observations of individual motor rotation have shown 26 steps per revolution^22,23^, and these data have been interpreted as the steps being generated by the repeated association and dissociation of the stator protein MotA and the rotor protein FliG for torque generation because the number of FliF-FliG subunits forming the MS ring and part of the C ring was thought to be 25 to 26 at the time^24–26^. However, recent structural analysis of the MS ring revealed its symmetry to be 33-fold with a variation from 32 to 35 (Ref.^27^), and the analysis of the flagellar basal body showed the MS ring to have 34-fold symmetry without variation and the C ring to have a small symmetry variation around 34-fold^28^. These results indicate the number of FliG on the C ring is mostly 34 and therefore invalidate the previous interpretation that the 26 steps per revolution reflect the elementary process of torque generation. Because the LP ring has 26-fold symmetry, the observed steps may be caused by potential minima formed by electrostatic force interactions between the rod and LP ring.

**Fig. 4.**
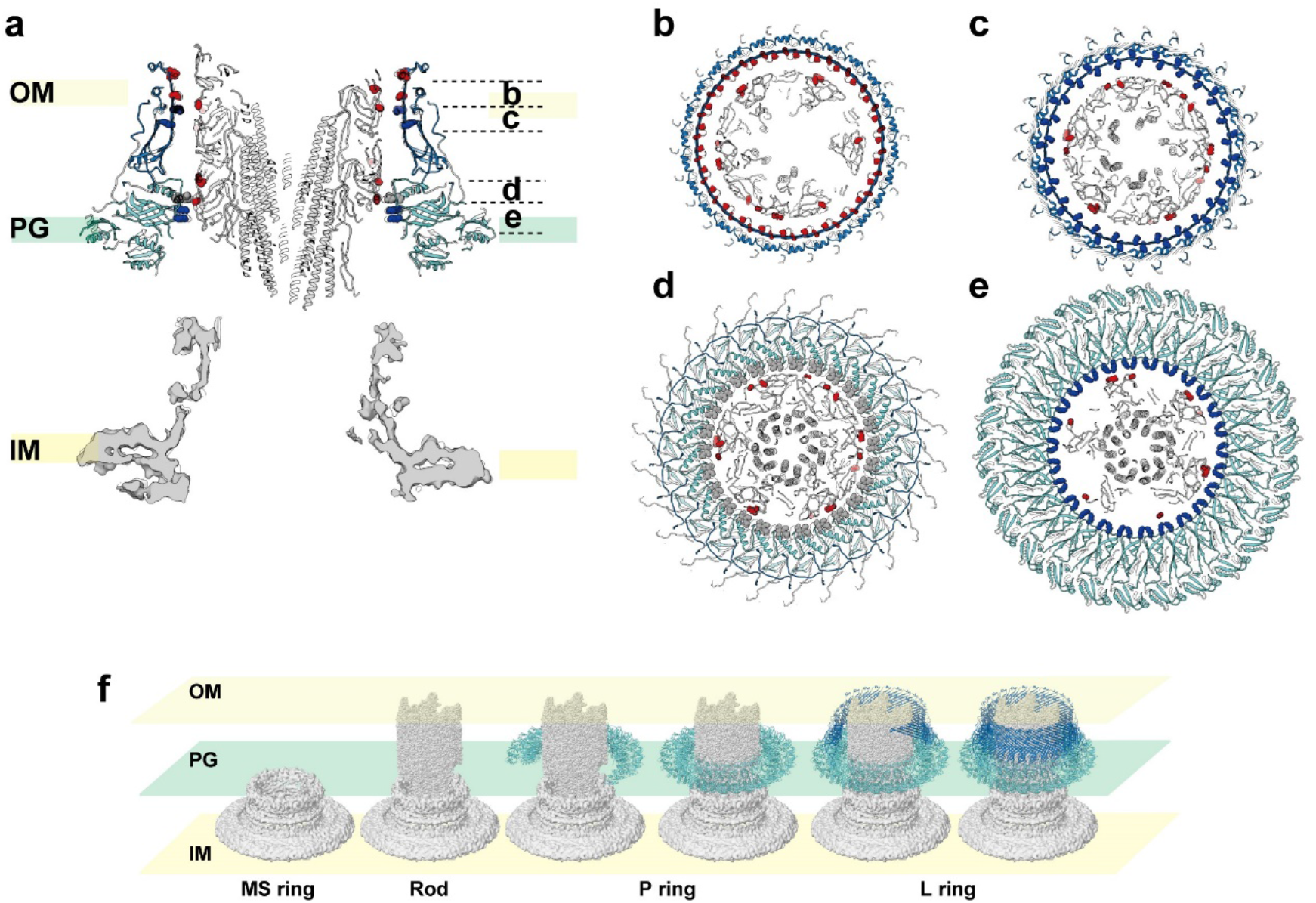
Interactions between the rod and LP ring and their assembly process. **a**, Vertical section of the atomic model of the rod (gray), L ring (sky-blue), P ring (cyan) and the map of the MS ring (gray). Charged amino acid residues are displayed with negative in red (Asp-109, Asp-154, Glu-203 of FlgG and Asp-78, Asp-86, Glu-104 of FlgH) and positive in blue (Lys-69, Lys-114 of FlgH and Lys-63, Lys-95 of FlgI). Four polar residues from Thr-33 to Thr-36 of FlgI in the P ring loop, which is located most closely to the rod surface, are shown in gray. **b**, **c**, **d**, **e**, Axial view of four horizontal slices of the rod and LP ring as indicated in **a**, showing repulsive and attractive interactions between the rod and LP ring: **b**, repulsive interactions; **c**, attractive interactions; **d**, the smallest gap between the rod and LP ring with non-charged residues of FlgI in the P ring loop; **e**, attractive interactions. **f**, The assembly process of the rod and LP ring.

Although the LP ring is a stable component of the hook-basal body even after its isolation from the cell, the LP ring easily slips off the rod when the basal body is isolated without the hook^6,7,28^, indicating very weak interactions between the rod and LP ring to keep the LP ring in the proper position around the rod. In the absence of FlgH, however, the basal body formation stops just after completion of P ring assembly by FlgI around the rod to form a structure called “candlestick”^7,29^, indicating that the P ring is tightly attached to the rod until the L ring is assembled above it. Among the *flgG* mutants that form the polyrod, many P rings are formed around the polyrod by *flgG* (G65V) and *flgG* (P52L) mutants whereas no P ring is observed by FlgG mutations of G53R, G183R or deletion of residues 54–57. All of these FlgG mutations are located in the long β-hairpin structure called “L-stretch” formed by *Salmonella* FlgG specific sequence not present in FlgE (Extended Data Fig. 8a,b), and its tip is presumably exposed on the surface of the rod^8,30^ (Extended Data Fig. 8b–e). Our model of the LP ring shows that Lys-63 and Lys-95 of FlgI are located on the inner surface of the P ring and directly face the negatively charged surface of the rod. Their mutations to either alanine to diminish the positive charge or aspartate to have the negative charge impair cell motility possibly by inhibiting P ring formation (Fig. 3b,c). So, these two lysine residues are likely to play important roles in P ring assembly on the negatively charged surface of the rod. This is in agreement with no P ring assembly by FlgG polyrod mutations G53R and G183R as well as deletion of restudies 54-57, which would put Glu-58 of FlgG at the tip of the L-stretch too far from Lys-63 and Lys-95 of FlgI (Extended Data Fig. 8c–e). We therefore suggest that Lys-63 and Lys-95 of FlgI are essential for the initial binding of FlgI to the rod surface to form the P ring tightly attached to the rod and that some conformational changes occur in the P ring upon binding of FlgH for L ring assembly to make the completed LP ring free from the rod for its free, high-speed rotation within the LP ring as a bushing (Fig. 4f). Structural analysis of the P rings stably attached to the polyrod is underway to clarify the mechanism of LP ring assembly.

## Methods

### Bacterial strains, media, and bacterial growth

Bacterial strains and plasmids used in this study are listed in Extended Data Table 1. For cryoEM structural analyses, a strain expressing only the hook-basal body and not the filament (HK1002) was used. The L-broth and soft tryptone agar plate were used as described previously^31^. The soft agar plate used for swarming assay contains 1% (w/v) Bacto tryptone, 0.5% (w/v) NaCl and 0.35 % (w/v) Bact agar. Ampicillin was added to LB at a final concentration of 100 μg/mL.

### DNA manipulation

All primers used in this study were listed in Extended Data Table 2. The *flgHI* genes were amplified by TaKaRa Ex Taq® DNA polymerase (TaKaRa Bio) using the chromosomal DNA of the *Salmonella* wild-type strain SJW1103 as a template and primers, FlgHI_NedI_Fw_2 and FlgHI_BamHI_Rv_3. The PCR products were digested with NdeI and BamHI and cloned into the NdeI and BamHI sites of the pET22b vector (Novagen) to generate a plasmid, pTY03. In order to clarify whether the positively charged Lys-63 and Lys-95 of FlgI is important for forming the P ring upon interacting with negatively charged surface of FlgG, each or both Lys-63 and Lys-95 were replaced by Ala or Asp residue. Site-directed mutagenesis was carried out using primer STAR® MAX DNA polymerase (TaKaRa Bio). The mutated genes were amplified by PCR using pTY03 as a template and pairs of complementary primers containing a mutagenized codon listed in Extended Data Table 2, and then the plasmids were introduced to *E. coli* NovaBlue cells (Novagen) for transformation.

### Hook-basal body (HBB) purification

The *Salmonella* HK1002 cells were pre-cultured in 30 mL LB with overnight shaking at 37°C and inoculated into a 2.6 L of fresh LB. The cells were grown until the optical density had reached an OD_600_ of about 1.0. The cells were collected by centrifugation and resuspend in 10% sucrose buffer [10% (w/v) sucrose, 0.1 M Tris-HCl, pH 8.0]. EDTA (pH 8.0) and lysozyme were added to final concentrations of 10 mM and 1.0 mg/mL, respectively. The cell lysates were stirred on ice for 1 hour at 4°C, and 0.1 M MgSO4 and 10% (w/v) Triton X-100 were then added to final concentrations of 10 mM and 1% (w/v), respectively. After stirring for 1 hour at 4°C, 0.1 M EDTA (pH 11.0) was added to a final concentration of 10 mM. The solution was centrifuged at 15,000 g, and the supernatant was collected. The pH was adjusted to 10.5 with 5 N NaOH and recentrifuged at 15,000 g to remove undissolved membrane fractions. The supernatant was centrifuged at 67,000 g to collect HBBs as a pellet. This pellet was resuspended in 1 mL of Buffer C [10 mM Tris-HCl, 5 mM EDTA, 1% (w/v) Triton X-100] and was centrifuged at 9,700 g, and the supernatant was collected. HBBs were purified by a sucrose density-gradient centrifugation method with a gradient of sucrose from 20% to 50%. Fractions containing HBBs were collected and checked by SDS-PAGE with Coomassie Brilliant Blue staining. The fractions containing HBBs were two-fold diluted by a final buffer [20 mM Tris-HCl, pH 8.0, 150 mM NaCl, 0.05% (w/v) Triton X-100], and the sucrose was removed by centrifugation at 120,000 g. Finally, HBBs were resuspended with 10-50 μL of the final buffer, and the sample solution was stored at 4°C.

### Electron cryomicroscopy and image processing

A 3 μL sample solution was applied to a Quantifoil holey carbon grid R1.2/1.3 Mo 200 mesh (Quantifoil Micro Tools GmbH, Großlöbichau, Germany) with pretreatment of a side of the grid by 10 sec glow discharge. The grids were plunged into liquid ethane at a temperature of liquid nitrogen for rapid freezing^32^ with Vitrobot Mark IV (Thermo Fisher Scientific) with a blotting time of 5 second at 4°C and 100% humidity. All the data collection was performed on a prototype of CRYO ARM 200 (JEOL, Japan) equipped with a thermal field-emission electron gun operated at 200 kV, an Ω-type energy filter with a 20 eV slit width and a K2 Summit direct electron detector camera (Gatan, USA). The 12,759 dose-fractionated movies were automatically collected using the JADAS software (JEOL). Using a minimum dose system, all movies were taken by a total exposure of 10 seconds, an electron dose of 0.9 electrons/Å^2^ per frame; a defocus range of 0.2 to 2.0 μm, and a nominal magnification of 40,000×, corresponding to an image pixel size of 1.45 Å. All the 50 frames of the movie were recorded at a frame rate of 0.2 sec/frame. The defocus of each image was estimated by Gctf-v1.06 (Ref.^33^) after motion correction by RELION-3.0-β2 (Ref.^34^). The HBB particle images were automatically picked by an in-house python program (YOLOPick) that utilizes a convolutional neural network program, YOLOv2 (Ref.^35^). In total, 64,418 particles were extracted into a box of 340 × 340 pixels and used for 2D classification by RELION 3.0-β2. First, clear 2D classes were selected (1,515 particles), and the “ignoring CTF first-peak” option was applied for the remaining blur 2D classes (62,841 particles) to select additional 15,902 particles, and in total, 17,363 particles were selected. The 15,242 particles were selected from those 17,363 particles after re-centering their positions by in-house program named TK-center.py, then the 14,975 particles were selected after another 2D classification, and those particles were used for first 3D classification into two classes with C1 symmetry. A density map of better quality, reconstructed from 13,017 particles (87%), was then used as a reference for density subtraction to leave only the LP ring. The C26 symmetry was visible in the LP ring density map, but its cross-section image was transformed from the Cartesian to the polar coordinates, and its autocorrelation function was Fourier transformed to confirm the rotational symmetry to be 26-fold. This estimation was carried out by in-house program named Tksymassign.py. Then, the C26 symmetry was applied for further image analysis to obtain a higher resolution 3D map, which resulted in 3.5 Å resolution at a Fourier shell correlation (FSC) of 0.143 after 3D refinement and CTF refinement procedure from 10,802 particles (EMD-30398) (Extended Data Figs. 1 and 2). In order to obtain detailed information on the relative positioning of the rod and LP ring, 14,975 HBB particle images were extracted into box of 400 × 400 pixels from the same movie data sets described above and 14,370 particles were selected after 3D classification and 3D refinement. Because the resolution was not improved by this 3D classification and 3D refinement, 2D classification and centering were also done to this data set. In order not to be affected by the hook density, the reference and mask without the hook density was used for next 3D classification and 3D refinement. A density map of 6.9 Å resolution (at the FSC of 0.143) was obtained with C1 symmetry after 3D refinement and CTF refinement procedure from 14,370 particles (EMD-30409).

### Model building

The atomic model of FlgH and FlgI was constructed by COOT (Crystallographic Object-Oriented Toolkit)^36^, and PHENIX-1.13-2998 was used for auto sharpening of the cryoEM LP ring map and real-space refinement^37^. The secondary structures were predicted by PSIPRED^38^. In order to identify the minimum units composing L ring and P ring, only the Cα atoms were placed on the cryoEM map by COOT to make the main chain model of FlgH and FlgI. For further refinement, the densities around 4 Å from the Cα atoms were extracted from the LP ring density map, and the model was automatically refined by PHENIX real-space refinement and manually fixed by COOT. The side chains were added manually by COOT based on the cryoEM map and PSIPRED prediction. In order to consider the interactions between adjacent molecules, seven FlgH and three FlgI molecules were used for further refinement, and the central models of FlgH and FlgI were extracted to be the final model. The entire LP ring model was made by UCSF Chimera^39^ using the C26 symmetry. The surface potentials were calculated by APBS^40^ and PDB2PQR^41^ and multi sequence alignment (conservation) was done by Clustal Omega^42^. All the figures of 3D maps and atomic models used in this paper were prepared by UCSF Chimera^39^.

### Motility assays

We transformed *Salmonella* SJW203 cells (Δ*flgH-flgI*) with a pET22b-based plasmid encoding FlgH and FlgI (pTY03) or its mutant variants. Fresh transformants are inoculated onto soft agar plates [1% (w/v) tryptone, 0.5% (w/v) NaCl, 0.35% Bacto agar] with 100 μg/mL ampicillin and incubated at 30°C. At least seven independent measurements were performed.

### Secretion assays

*Salmonella* SJW203 cells (Δ*flgH-flgI*) harboring a pET22b-based plasmid encoding FlgH and FlgI (pTY03) or its mutant variants were grown overnight in 5 mL L-broth with gentle shaking at 30ºC. Details of sample preparations have been described previously^43^. Both whole cellular proteins and culture supernatants were normalized to a cell density of each culture to give a constant cell number. After sodium dodecyl sulfate-polyacrylamide gel electrophoresis (SDS-PAGE), immunoblotting was carried out with polyclonal anti-FlgE, anti-FliC, anti-FlgH or anti-FlgI antibody^21^. Detection was performed with an ECL prime immunoblotting detection kit (GE Healthcare). Chemiluminescence signals were captured by a luminoimage analyzer LAS-3000 (GE Healthcare). At least three independent measurements were performed.

### Reporting summary

Further information on research design is available in the Nature Research Reporting Summary linked to this paper.

### Data availability

Density maps are available at EMDB with accession codes EMD-30409 (HBB) and EMD-30398 (LP ring). Atomic models are available in Protein Data Bank with accession codes 7CLR (LP ring). Other data supporting this study are available from the corresponding authors on reasonable request.

## Author information

### Contributions

T.K. and K.N. designed the project; F.M, T.K. T.Minamino, and K.N. designed the experiments; T.Y., F.M., T.Miyata, and T.Minamino prepared the samples; T.K. set up the cryoEM imaging and analysis system; T.Y., F.M., T.Miyata, T.K. collected and analyzed cryoEM image data and built the atomic model; T.Y. and T.Minamino performed genetic, biochemical and physiological experiments; all authors studied the atomic model; T.Y. and K.N. wrote a draft of the paper, and all the other authors discussed and contributed to writing up the paper.

## Acknowledgments

We thank Kelly Hughes for providing strain SJW203, Hideyuki Matsunami for the HK1002 strain, Miki Kinoshita for help in genetic engineering and biochemical assay and for providing plasmids and Katsumi Imada for help and advice in constructing the atomic model.

