## Supplemental Materials for "Structure of the molecular bushing of the bacterial flagellar motor"

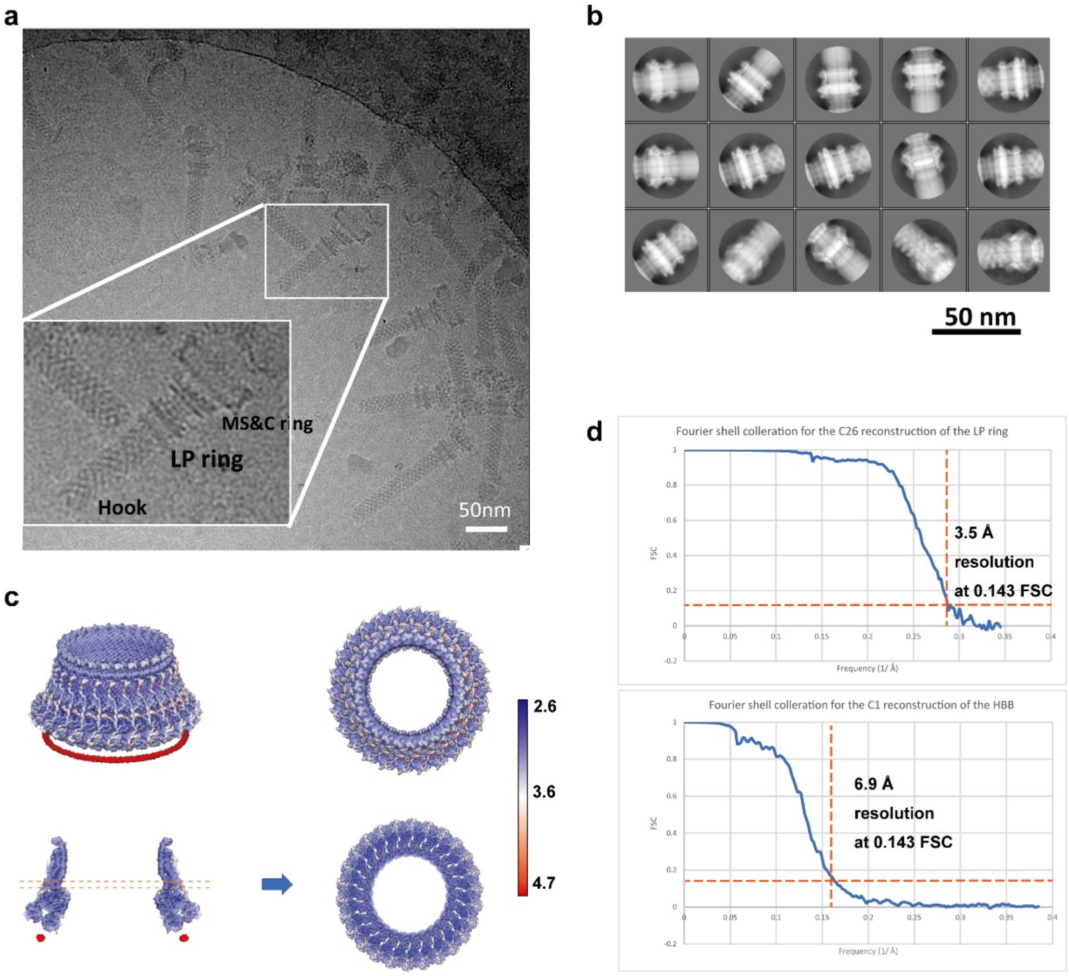

**Extended Data Fig. 1 CryoEM single particle image analysis of the LP ring**. **a**, Representative micrograph of the HBB complex. **b**, Selected 2D class averages used for 3D reconstruction of the LP ring with C26 symmetry. **c**, The local resolution of the final C26 LP ring density map colored from red (4.7 Å) to blue (2.6 Å). **d**, FSCs of the final density map of the LP ring reconstructed with C26 symmetry (upper) and the HBB complex with C1 symmetry (lower).

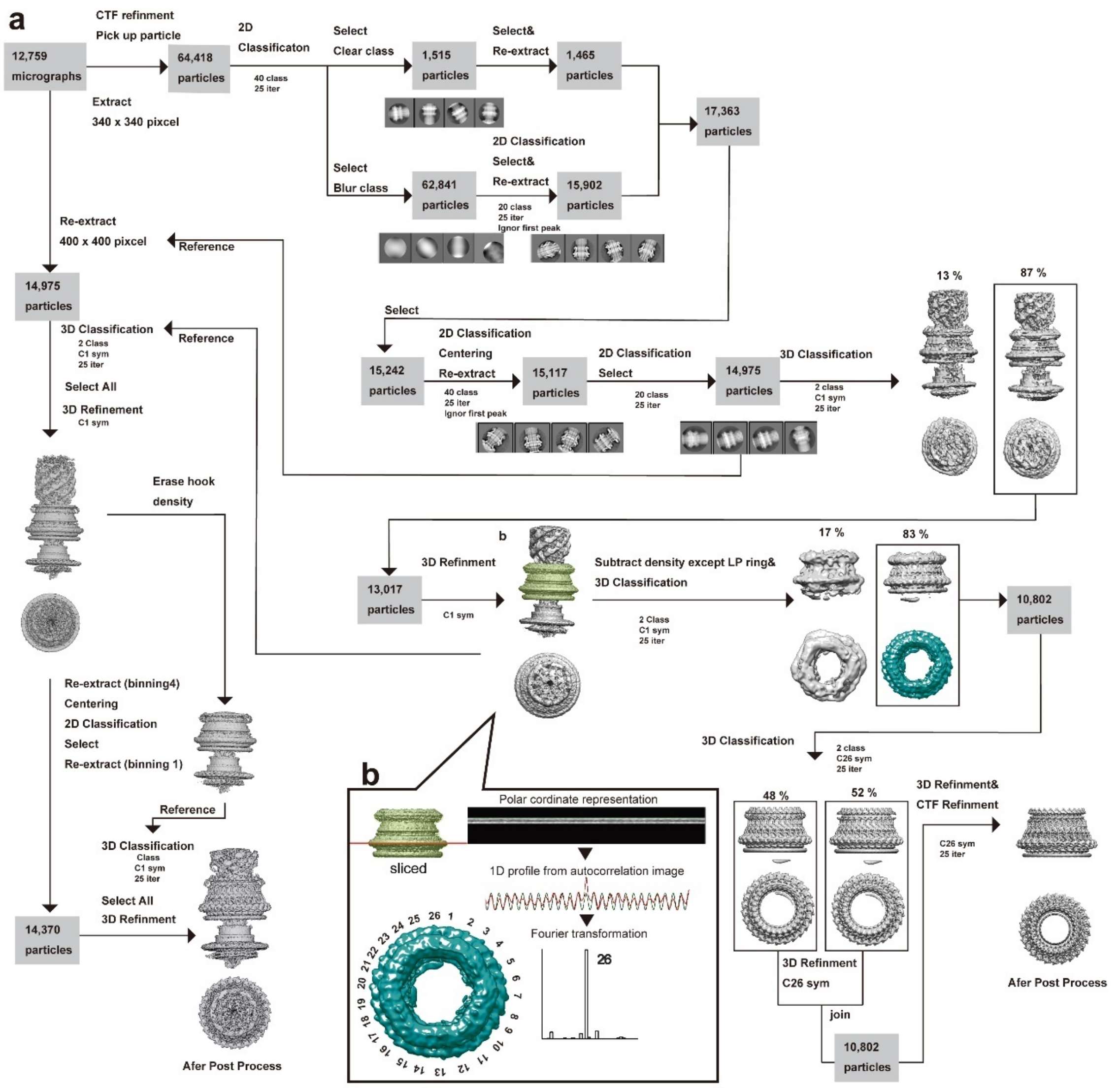

**Extended Data Fig. 2 Work process of cryoEM 3D reconstruction of the HBB complex and LP ring. a**, The work process of cryoEM single particle structural analysis of the HBB complex, in which 14,370 particles were used for final 3D classification and 3D refinement to produce a HBB density map at 6.9 Å resolution, and that of the LP ring alone, in which 10,802 particles were used to produce a density map at 3.5 Å resolution. **b**, The LP ring density colored light green was used to analyze its rotational symmetry. A horizontal slice of the LP ring density map at the red line was converted into the polar coordinate, and its auto-correlation was calculated and Fourier transformed to confirm the C26 symmetry indicated by visual inspection of the top view of the LP ring (green) extracted from the HBB complex reconstructed with C1 symmetry.

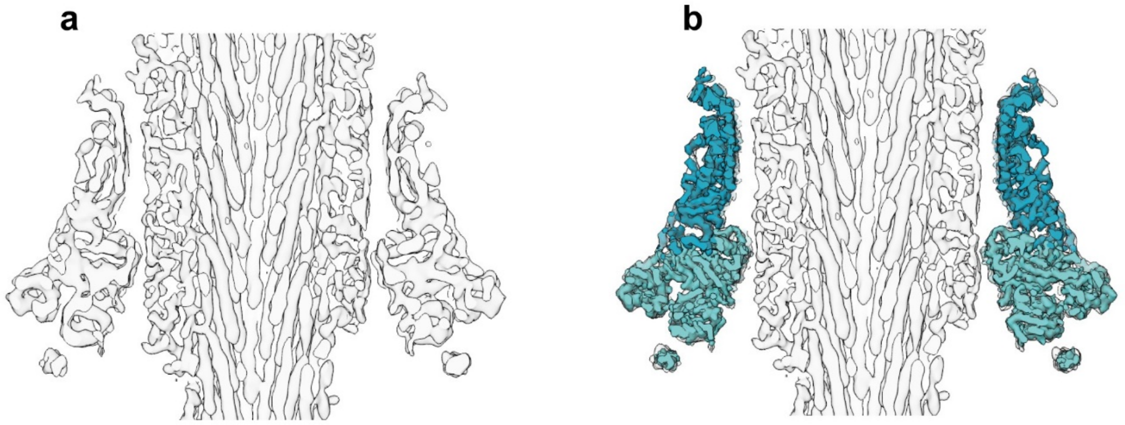

**Extended Data Fig. 3 The LP ring position relative to the rod determined from the HBB complex density map. a**, Vertical slice of the HBB complex density map (C1) including the LP ring and distal rod reconstructed at 6.9 Å resolution. **b**, The LP ring density map at 3.5 Å resolution (C26) superimposed on the figure in **a**.

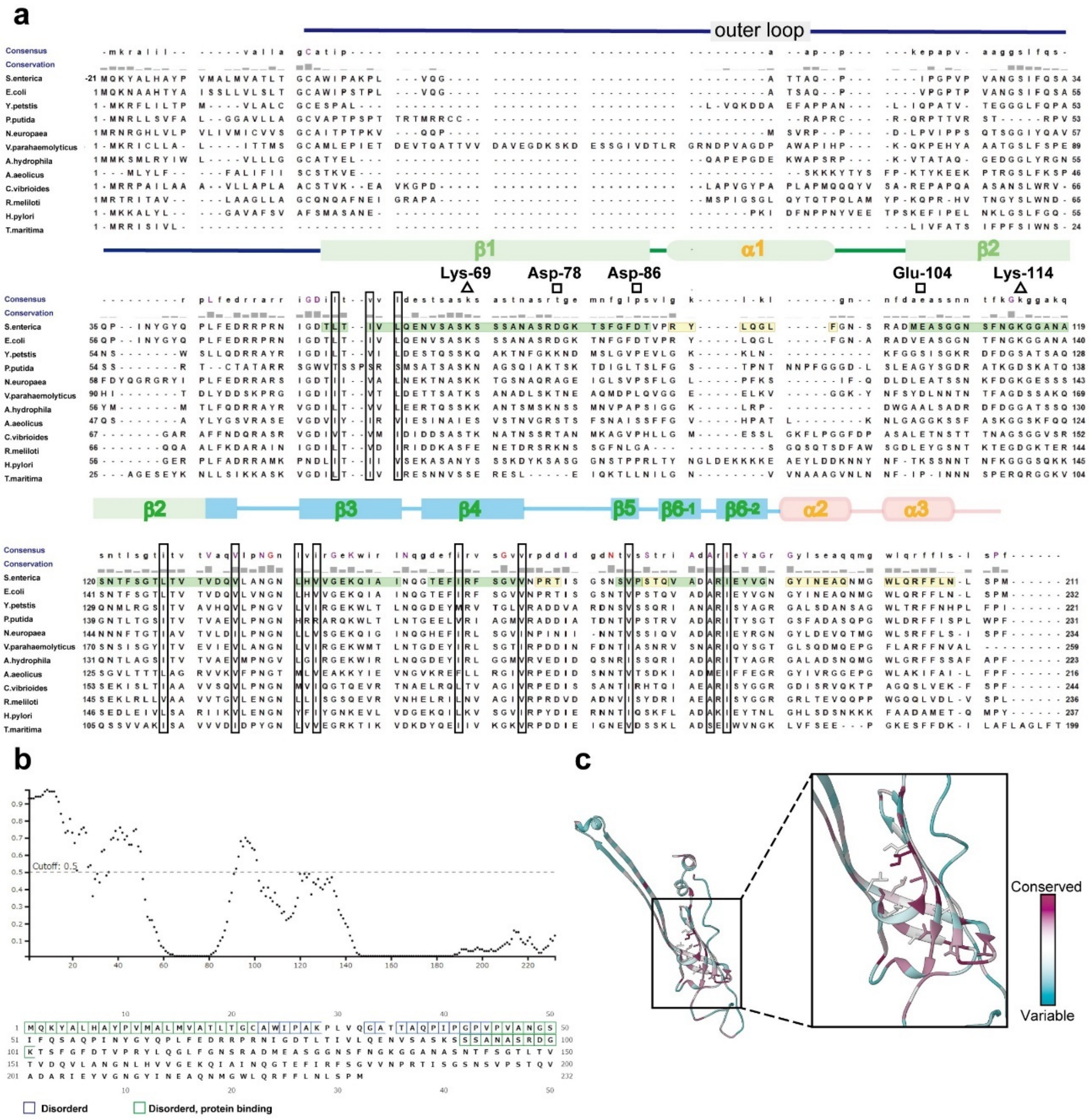

**Extended Data Fig. 4 Multiple sequence alignment of FlgH.** **a**, Multiple sequence alignment of FlgH carried out by Clustal Omega[42]. UniProt Accession numbers: *S. enterica* serovar Typhimurium (here after *S. enterica*), P0A1N8; *E. Coli*, P0A6S0; *Y. pestis*, Q8ZHZ5; *P. putida*, Q52081; *N. europaea*, Q82XG6; *V. parahaemolyticus*, Q9X9J5; *A. hydrophila*, A0A142DZW5; *A. aeolicus*, O67609; *C. vibrioides*, A0A0H3C9Y8; *R. meliloti*, A0A222JND2; *H. pylori*, E8QV65; *T. maritima*, Q9X1M6. The five charged residues (Lys-69, Asp-78, Asp-86, Glu-104, Lys-114) and color-coded secondary structures of *Salmonella* FlgH in three domains (outer layer, blue; inner layer, light green; middle layer, cyan) are shown above the sequence. **b**, Prediction of disordered regions calculated by PSIPRED[38]. The disorder plot (upper) and predicted disordered regions (lower) agree with the outer layer chain of our atomic model. **c**, the sequence conservation of FlgH is colored from cyan to magenta. Side chains involved in intramolecular hydrophobic interactions are shown in stick representation and magnified (right), and the hydrophobic residues are also boxed in **a**.

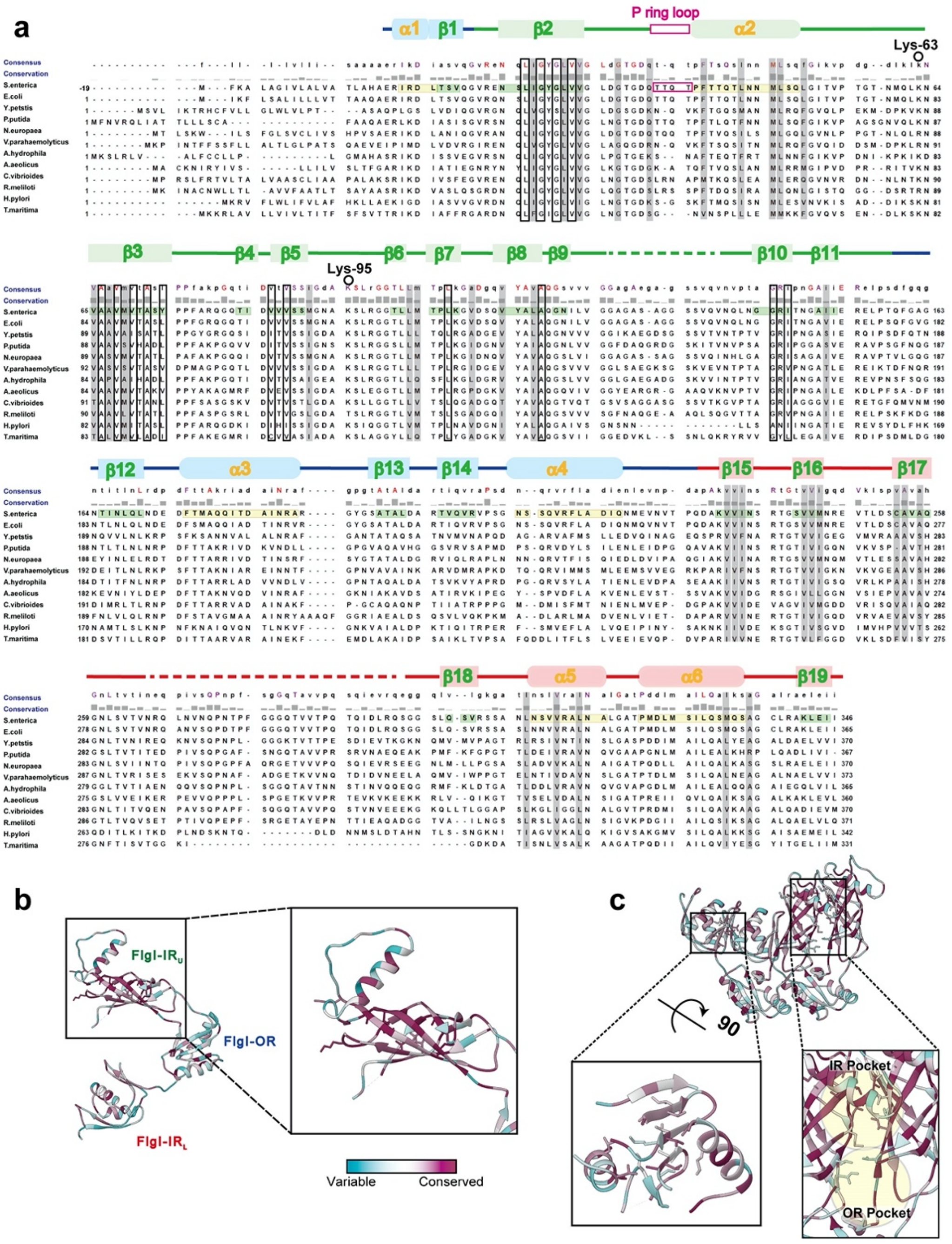

**Extended Data Fig. 5 Multiple sequence alignment for FlgI. a**, Multiple sequence alignment of FlgI carried out by Clustal Omega[42]. UniProt Accession numbers: *S. enterica*, P15930; *E. Coli*, P0A6S3; *Y. pestis,* Q8ZHZ4; *P. putida,* Q52082; *N. europaea,* Q82XG5; *V. parahaemolyticus*, Q87JI2; *A. hydrophila,* A0KM38; *A. aeolicus,* O67608; *C. vibrioides,* P33979; *R. meliloti,* Q52948; *H. pylori,* O25028; *T. maritima*, Q9X1M5. The two charged residues (Lys-63, Lys-95) and color-coded secondary structures of *Salmonella* FlgI with three domains (FlgI-OR, cyan; FlgI-IR$_{U}$, light green; FlgI-IR$_{L}$, red) are shown above the sequence. **b**, **c**, The sequence conservation of FlgI is colored from cyan to magenta. Side chains involved in intra and intermolecular hydrophobic interactions are shown in stick representation and magnified in **b** and **c**, respectively, and the hydrophobic residues are also boxed and gray meshed in **a**, respectively.

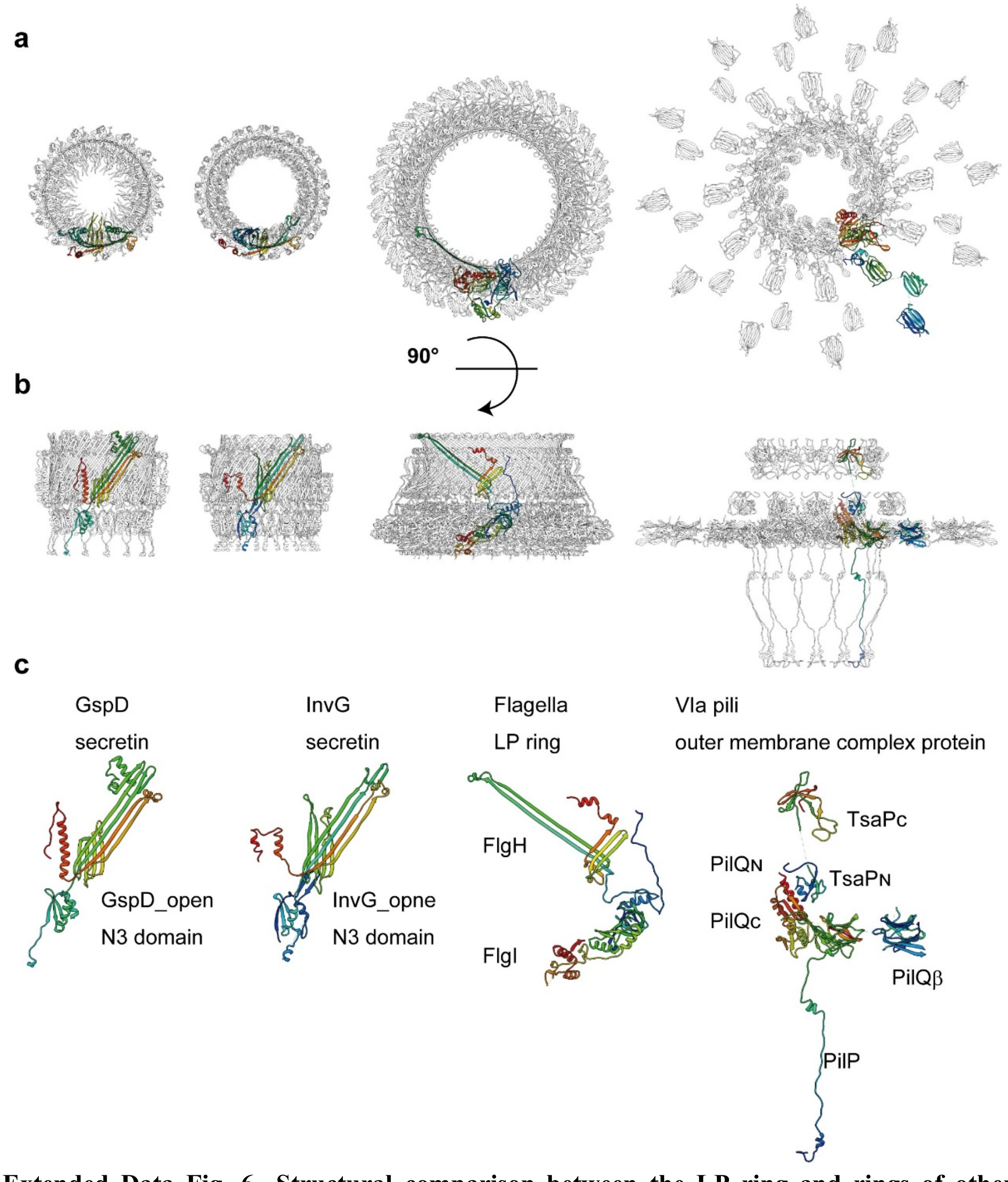

**Extended Data Fig. 6 Structural comparison between the LP ring and rings of other secretion systems. a**, **b**, Top and side views of GspD_open (residues 232–617, PDB ID: 5WQ7), InvG_open (residues 176–227, 252-557, PDB ID: 6DV6), the LP ring (PDB ID: 7CLR) and PilO, P, Q and TsaP complex of VIa pili (PDB ID: 3JC9). **c,** Side view of each monomer colored rainbow.

**a**

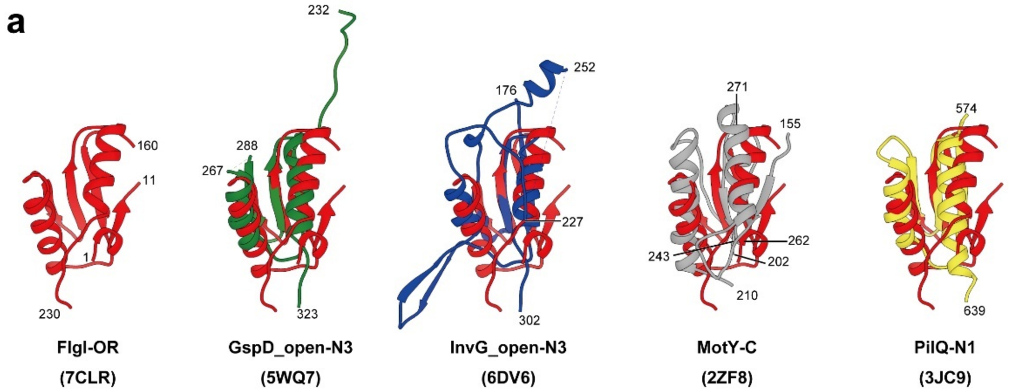

**b**

| | GspD_open-N3 | IngG_open-N3 | MotY-C | PilQ-N1 |
|---|---|---|---|---|
| [RMS(Å)] root mean square deviation for aligned Ca atoms | 3.561 | 2.694 | 0.708 | 3.628 |
| [DRMS(Å)] root mean square deviation for aligned Cb atoms | 3.210 | 2.293 | 0.907 | 3.194 |
| [SqID(%)] sequence identity | 11.3 | 10.3 | 7.70 | 6.40 |

**c**

| | PDB | Sequence |
|---|---|---|
| FlgI-OR | 7CLR | 1-11,160-230 |
| GspD_open-N3 | 5WQ7 | 232-267, 288-323 |
| InvG_open-N3 | 6DV6 | 176-227, 252-302 |
| MotY-C | 2ZF8 | 155-202, 210-243, 262-271 |
| PilQ-N1 | 3JC9 | 574-639 |

**d**

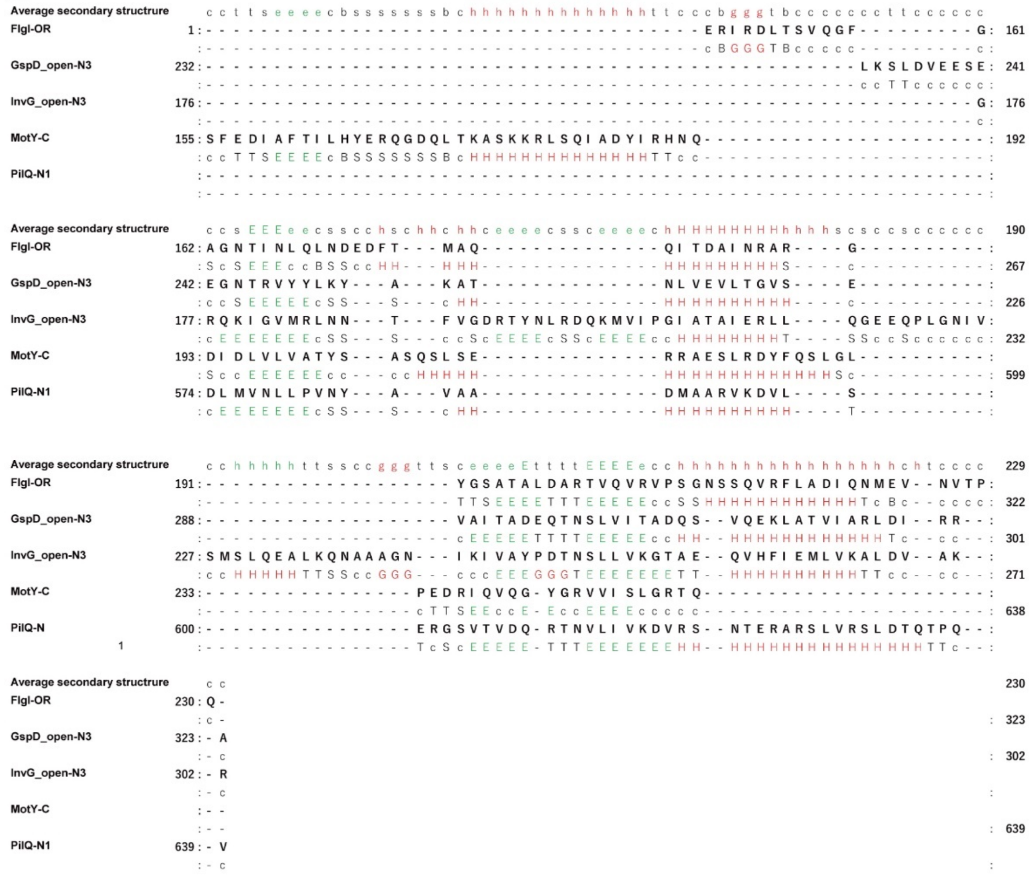

**Extended Data Fig. 7 Structural comparison between FlgI-OR and related domains of other secretion systems. a**, Superposition of FlgI-OR (residues 1–11, 16–230: red) on to GspD_open-N3 (residues 232–267, 288-323: green), InvG_open-N3 (residues 176–227, 252-302: blue), MotY-C (PDB ID: 2ZF8, residue 155-202, 210-243, 262-271: gray) and PilQ-N1 (residues 574–639: yellow). **b**, Alignment score between FlgI-OR and other four domains. **c**, Color , Name, PDB ID, and sequence region of compared domains. **d**, Sequence alignment between FlgI-OR and other four domains. All alignment and tables were calculated by MATRAS[20].

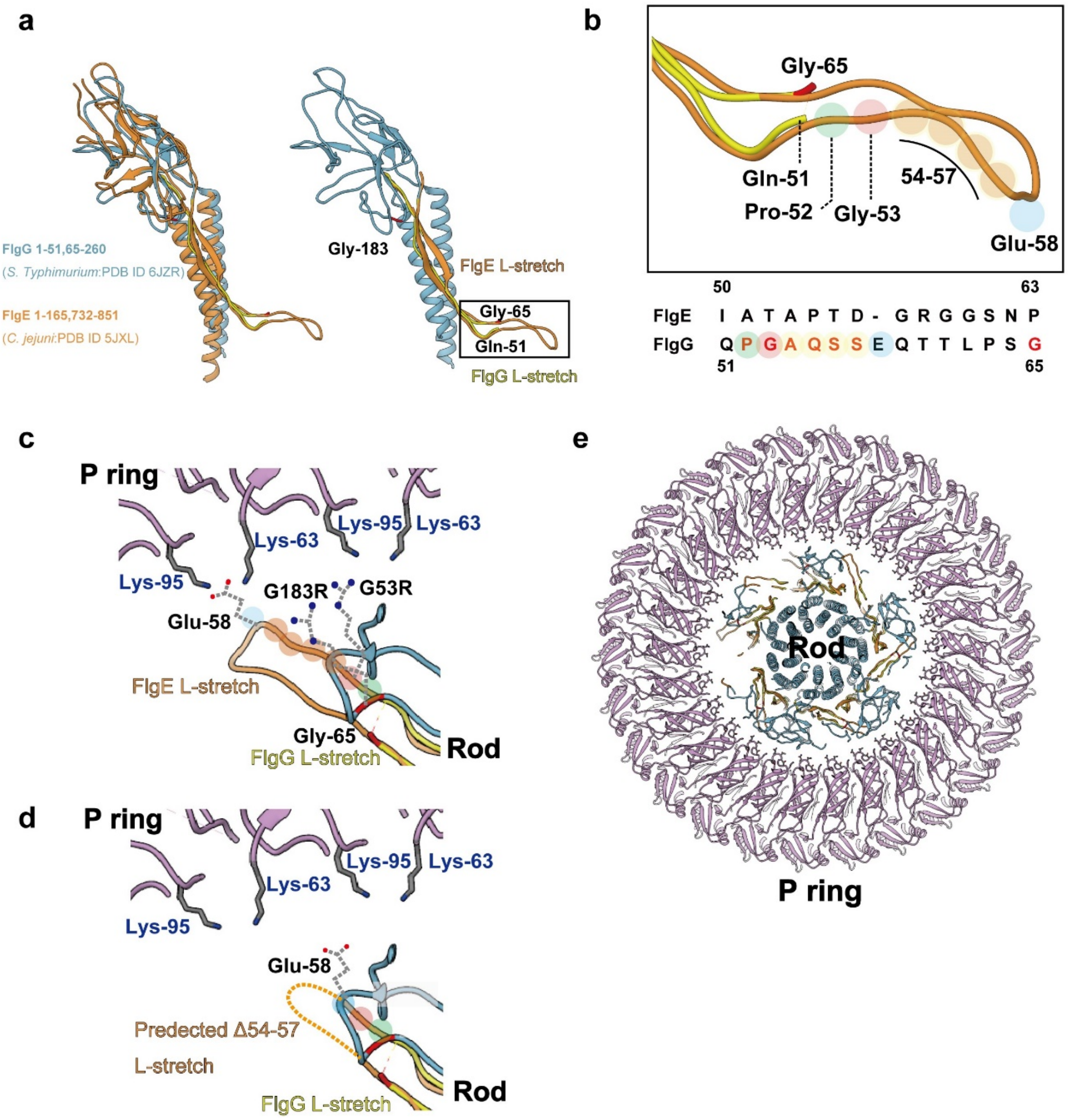

**Extended Data Fig. 8 Interactions between FlgG of the rod and FlgI of the P ring. a**, Structural comparison between FlgG of *S. enterica* (PDB ID: 6JZR) and FlgE (residues 1-165 and 732-851) of *C. jejuni* (PDB ID: 5JXL). The FlgE L-stretch (residues 31–80) is superimposed to FlgG L-stretch (residues 32–82) with RMSD 1.16 Å. **b**, The position of the missing residues of the FlgG model (residues 52–58) in the L-stretch, with residues predicted to interact with FlgI in the P ring are displayed by circles. **c**, Possible interactions of Lys-63 and Lys-95 of FlgI in the P ring and Glu-58, Arg-183 and Arg-53 of FlgG in the rod. Glu-58 is expected to interact with Lys-63 and Lys-95 and help P ring assembly. The G183R and G53R mutation of FlgG is expected to disturb P ring assembly by repulsive force. **d**, Expected change of the Glu-58 position in the Δ54-57 mutant of FlgG, making the distance between Glu-58 and Lys-63 and Lys-95 of FlgI too far for interaction. **e**, Horizontal slice of the rod and P ring at their closest position, with side chains of FlgI in the P ring loop are displayed in stick representation. There is still a small gap between the rod and P ring to accommodate water molecules.

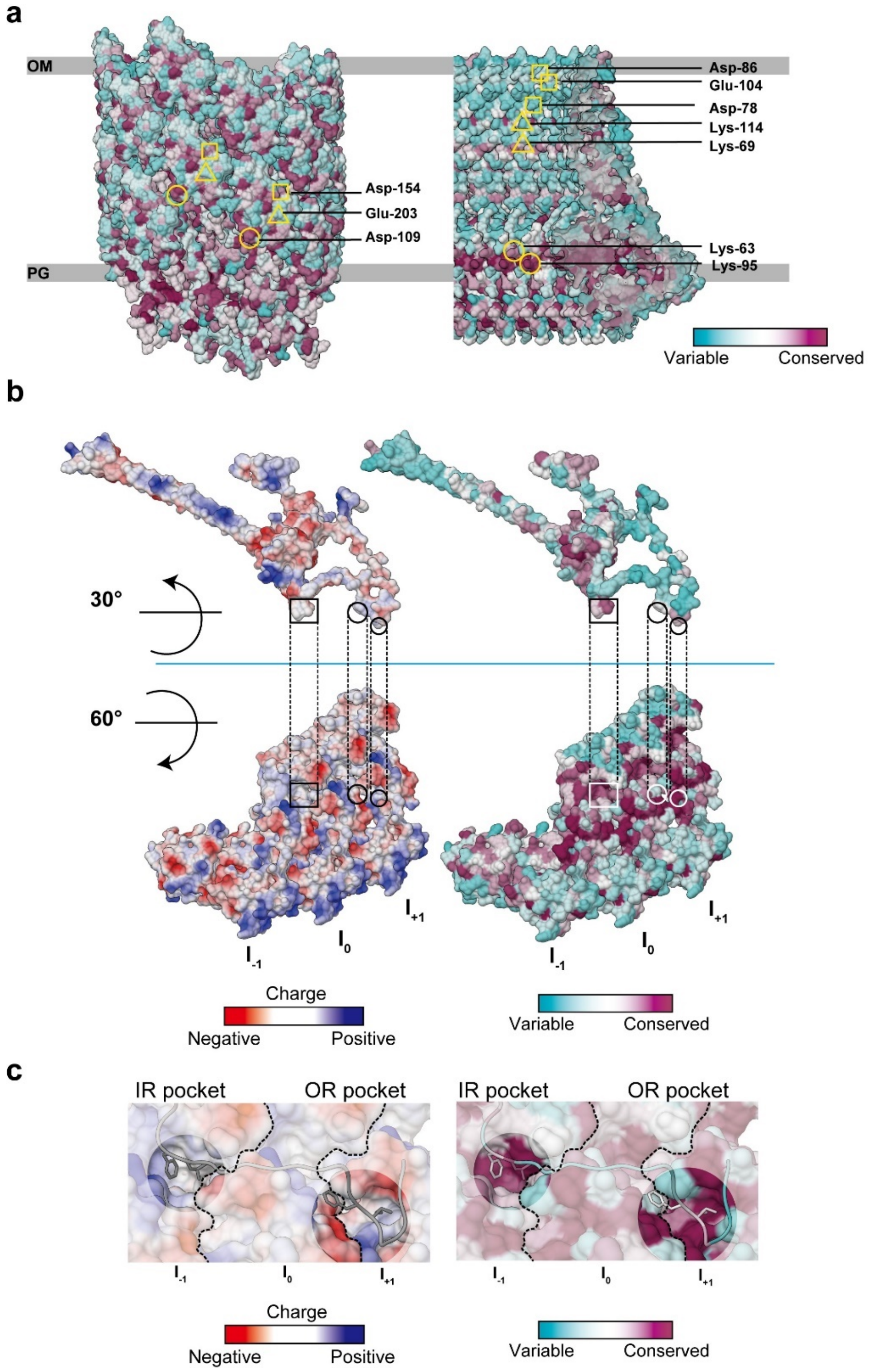

**Extended Data Fig. 9 Conservation and electrostatic surface potential of the rod, FlgH, FlgI.** **a**, Conservation on the outer surface of the distal rod and the inner surface the LP ring. Asp-109 (○) and Aps-154 (□) of the rod and Lys-63 and Lys-95 (○) of the P ring are highly conserved. **b**, Interactions between one FlgH and three FlgI ($I_0$, $I_{\pm1}$) molecules. The interactions in the IR (□) and OR (○) pockets are both hydrophobic (left). The residues consist of the IR pocket are highly conserved (right). **c**, Enlarged views of the interactions in the IR and OR pockets. The three FlgI ($I_0$, $I_{\pm1}$) molecules are displayed as solid surface and FlgH in Cα ribbon. The hydrophobic side chains of FlgH are displayed in stick and are color-coded as gray in the electrostatic surface potential (left) and cyan to magenta in the conservation (right).

**Extended Data Table 1  Strains and plasmids used in this study**

| Strains and plasmid | Relevant characteristics | Source or reference |
|---|---|---|
| ***Salmonella* strains** | | |
| HK1002 | Not forming the filament<br>Δ*flgK*, Δ*cliP::*$Cm^r$ | Ref.[44] |
| SJW203 | LP ring deletion mutant<br>Δ*flgH-flgI* | Shigeru Yamaguchi & Ref.[45] |
| **Plasmids** | | |
| pET22b | Expression vector | Novagen |
| pTY03 | pET22b/FlgH+FlgI | This study |
| pTY03(K63A) | pET22b/FlgH+FlgI(K63A) | This study |
| pTY03(K63D) | pET22b/FlgH+FlgI(K63D) | This study |
| pTY03(K95A) | pET22b/FlgH+FlgI(K95A) | This study |
| pTY03(K95D) | pET22b/FlgH+FlgI(K95D) | This study |
| pTY03(K63A/K95A) | pET22b/FlgH+FlgI(K63A/K95A) | This study |
| pTY03(K63A/K95D) | pET22b/FlgH+FlgI(K63A/K95D) | This study |
| pTY03(K63D/K95A) | pET22b/FlgH+FlgI(K63D/K95A) | This study |
| pTY03(K63D/K95D) | pET22b/FlgH+FlgI(K63D/K95D) | This study |

**Extended Data Table 2  Primers used in this study**

| Mutated residue | Primer | Sequence (5'-3') |
|---|---|---|
| | FlgHI_NdeI_Fw-2 | CATATGCAAAAATACGCGCTTCACGC |
| | FlgHI_BamHI_Rv-3 | CTCGGATCCTCAGATGATTTCCAGTTTGGCGCG |
| K63A | FlgI_K63A_Fw | CAGTTGGCAAACGTGGCGGCGGTGATGGTGACG |
| | FlgI_K63A_Rv | CACGTTTGCCAACTGCATATTGGTGCCGGTGGG |
| K63D | FlgI_K63D_Fw | CAGTTGGATAACGTGGCGGCGGTGATGGTGACG |
| | FlgI_K63D_Rv | CACGTTATCCAACTGCATATTGGTGCCGGTGGG |
| K95A | FlgI_K95A_Fw | AACGCTGCAAGTCTGCGTGGCGGGACGTTATTA |
| | FlgI_K95A_Rv | CAGACTTGCAGCGTTCCCCATTGAGGAAACAAC |
| K95D | FlgI_K95D_Fw | AACGCTGATAGTCTGCGTGGCGGGACGTTATTA |
| | FlgI_K95D_Rv | CAGACTATCAGCGTTCCCCATTGAGGAAACAACGAC |
| K63A/ K95A | FlgI_K63A_Fw | CAGTTGGCAAACGTGGCGGCGGTGATGGTGACG |
| | FlgI_K63A_Rv | CACGTTTGCCAACTGCATATTGGTGCCGGTGGG |
| | FlgI_K95A_Fw | AACGCTGCAAGTCTGCGTGGCGGGACGTTATTA |
| | FlgI_K95A_Rv | CAGACTTGCAGCGTTCCCCATTGAGGAAACAAC |
| K63A/K95D | FlgI_K63A_Fw | CAGTTGGCAAACGTGGCGGCGGTGATGGTGACG |
| | FlgI_K63A_Rv | CACGTTTGCCAACTGCATATTGGTGCCGGTGGG |
| | FlgI_K95D_Fw | AACGCTGATAGTCTGCGTGGCGGGACGTTATTA |
| | FlgI_K95D_Rv | CAGACTATCAGCGTTCCCCATTGAGGAAACAACGAC |
| K63D/K95A | FlgI_K63D_Fw | CAGTTGGATAACGTGGCGGCGGTGATGGTGACG |
| | FlgI_K63D_Rv | CACGTTATCCAACTGCATATTGGTGCCGGTGGG |
| | FlgI_K95A_Fw | AACGCTGCAAGTCTGCGTGGCGGGACGTTATTA |
| | FlgI_K95A_Rv | CAGACTTGCAGCGTTCCCCATTGAGGAAACAAC |
| K63D/K95D | FlgI_K63D_Fw | CAGTTGGATAACGTGGCGGCGGTGATGGTGACG |
| | FlgI_K63D_Rv | CACGTTATCCAACTGCATATTGGTGCCGGTGGG |
| | FlgI_K95D_Fw | AACGCTGATAGTCTGCGTGGCGGGACGTTATTA |
| | FlgI_K95D_Rv | CAGACTATCAGCGTTCCCCATTGAGGAAACAACGAC |

**Extended Data Table 3 Summary of cryoEM data collection and refinement statistics**

| Data collection and processing | | |
|---|---|---|
| | HBB (EMDB-30409) | LP ring (EMD-30398) |
| CryoTEM | CRYO ARM 200 (prototype) | |
| Voltage (kV) | 200 | |
| Camera | K2 Summit | |
| Magnification | 40,000 | |
| Pixel size (Å / pix) | 1.45 | |
| Total exposure time (sec) | 10 | |
| No. of frames | 50 | |
| Frame rate (sec / frame) | 0.2 | |
| Dose rate par frames ($e^-$ / Å) | 0.9 | |
| Defocus range (μm) | 0.2 - 2.0 | |
| No. of micrographs | 12,759 | |
| No. of final particle images | 14,370 | 10,802 |
| Symmetry | C1 | C26 |
| Resolution (Å) | 6.9 | 3.5 |
| FSC threshold | 0.143 | |
